# Comparison of the Hi-C, GAM and SPRITE methods by use of polymer models of chromatin

**DOI:** 10.1101/2020.04.24.059915

**Authors:** Luca Fiorillo, Francesco Musella, Rieke Kempfer, Andrea M. Chiariello, Simona Bianco, Alexander Kukalev, Ibai Irastorza-Azcarate, Andrea Esposito, Mattia Conte, Antonella Prisco, Ana Pombo, Mario Nicodemi

## Abstract

Powerful technologies have been developed to probe chromatin 3D physical interactions genome-wide, such as Hi-C, GAM and SPRITE. Due to their intrinsic differences and without a benchmarking reference, it is currently difficult to assess how well each method represents the genome 3D structure and their relative performance. Here, we develop a computational approach to implement Hi-C, GAM and SPRITE *in-silico* to compare the three methods in a simplified, yet controlled framework against known polymer 3D structures. We test our approach on models of three 6-Mb genomic regions, around the *Sox9* and the *HoxD* genes in mouse ES cells, and around the *Epha4* gene in mouse CHLX-12 cells. The model-derived contact matrices consistently match Hi-C, GAM and SPRITE experiments. We show that *in-silico* Hi-C, GAM and SPRITE average data are overall faithful to the 3D structures of the polymer models. We find that the inherent variability of model single-molecule 3D conformations and experimental efficiency differently affect the contact data of the different methods. Similarly, the noise-to-signal levels vary with genomic distance differently in *in-silico* Hi-C, SPRITE and GAM. We benchmark the performance of each technology in bulk and in single-cell experiments, and identify the minimal number of cells required for replicates to return statistically consistent chromatin contact measures. Under the same experimental conditions, SPRITE requires the lowest number of cells, Hi-C is close to SPRITE, while GAM is the most reproducible method to capture interactions at large genomic distances.

## INTRODUCTION

The 3-dimensional (3D) organization of chromosomes in the nucleus of cells has a crucial role in the regulation of genomic activities and transcriptional programs^1–7^. To access genome-wide information on chromatin architecture and DNA interactions, a number of sequencing approaches are currently being developed^8^, while high-resolution microscopy is rapidly advancing^9–12^. Sequencing approaches include *3C*-based methods, such as Hi-C and its developments^13–19^, Genome Architecture Mapping (GAM)^20^ and Split-Pool Recognition of Interactions by Tag Extension (SPRITE)^21^. These approaches have shown that the mammalian genome has a complex 3D organization where functional contacts occur across distal DNA regions, such as loops between enhancers and promoters^15^, along with interactions at the megabase scale within Topologically Associated Domains (TADs)^22,23^ and higher-order structures such as metaTADs^24^ and A/B compartments^13^. However, it remains unclear to what extent these technologies are faithful to the underlying 3D structure of the genome and whether they measure different aspects of chromosomal 3D organization. Since they return distinct measures of chromatin interactions, it is also difficult to identify a clear benchmark to compare their performances in different conditions.

Hi-C methods have revolutionised the field of chromosome architecture and are widely used. They provide a measure of the abundance of pairwise interactions, i.e., a *Hi-C contact frequency map*, by sequencing the ligation products of DNA fragments that are in close spatial proximity in the nucleus^13,15^. GAM probes 3D proximity of DNA sites by sequencing the genomic content of thin cryo-sectioned and laser micro-dissected slices from the nuclei of cells fixed in optimal preservation conditions^8,20^. Physically distant DNA sites are unlikely to co-segregate in the same thin slice, whereas physically proximal sites tend to co-segregate. The output of a GAM experiment is a segregation table indicating which loci are present in each slice, based on sequencing its DNA content. To identify pair-wise or multi-ways contacts, a GAM *co-segregation map* is calculated, i.e., the frequency with which pairs (or groups) of genomic regions are found in the same slices. From GAM data single-cell DNA non-random interaction probabilities can be reconstructed by use of statistical tools, such as SLICE^20^. Finally, SPRITE^21^ relies on the sequencing of barcoded DNA: after DNA crosslinking and fragmentation in isolated nuclei (as in Hi-C), interacting chromatin complexes are uniquely barcoded via a split-pool method and identified by sequencing. SPRITE interaction maps can be extracted from analysing the DNA segments that have the same barcode, which must originate from the same interacting complex.

To compare Hi-C, GAM and SPRITE, we run a computational experiment that implements the three methods *in-silico* on an ensemble of known 3D polymer structures and analyse their outputs in a simplified, yet fully controlled framework. To facilitate the comparison with real experimental data, rather than using arbitrary polymer conformations, we focused on the polymer models of two genomic regions around the *Sox9* and the *HoxD* genes from mouse embryonic stem cells (mESC), respectively 6Mb and 7Mb long^25,26^. We also considered the model of a 6Mb long region around the *Epha4* gene from mouse CHLX-12 cells^27^. The comparison of the performance of the different technologies in those loci is interesting also because, for instance, disease-linked structural variants located around the *Sox9* and *Epha4* genes have been shown to induce gene mis-expression as a consequence of the rewiring of contacts with local enhancers^6,27,28^; and the *HoxD* locus presents a complex 3D compartmentalization which is thought to have a broad functional role in controlling transcriptional states during differentiation^29,30^. Different computational approaches^31–34^ and polymer models^25,27,35–44^ have been discussed to reconstruct chromatin 3D conformations. Here, we focused on the *String&Binders* (SBS) polymer model^27,37,39^ because it has been already validated against Hi-C data in those loci^25–27^. The polymer models of the considered loci were inferred from Hi-C data and used to derive an ensemble of known 3D structures for each locus. Those 3D structures were in turn employed to benchmark the performances of Hi-C, SPRITE and GAM in bulk as well as in single-cell computational experiments. For the *Sox9* locus, a polymer model inferred from GAM data has been also analysed^45^ and returned similar results. Finally, as a control, we also ran investigations in a toy block-copolymer model, unrelated to any chromosomal region, which returned a similar scenario about the performance of the three technologies.

Here, we show that *in-silico* average Hi-C, GAM and SPRITE matrices match their corresponding experimental bulk data validating our approach and the polymer models chosen. We found that bulk *in-silico* Hi-C, GAM and SPRITE data are all faithful to the reference 3D architecture as they have high correlations with the benchmark average distance matrix of our polymer structures. In contrast, single-cell data are affected by high levels of variability even in the case of ideal detection efficiency because of the inherent variety of single-molecule conformations of the polymer ensembles. Finally, we show that detection efficiency and number of single-molecule structures (a proxy for single cells) considered in the *in-silico* experiments differently affect contact data and the noise-to-signal ratio at different genomic distances across the three technologies. While poor efficiencies can be compensated by large cell numbers, we found that GAM is significantly less noise affected at larger genomic distances than the other methods, under similar experimental conditions (e.g., a given detection efficiency), but requires a comparatively much larger number of cells to ensure similarity across replicates, SPRITE requiring the least and Hi-C close to SPRITE.

## RESULTS

### Derivation of *in-silico* Hi-C, SPRITE, and GAM interaction maps from known single-molecule 3D structures

To compare *in-silico* Hi-C, GAM and SPRITE data, we focused first on the case study of a 6Mb-wide region around the *Sox9* gene (chr11:109Mb-115Mb, mm9) in mESCs. The SBS polymer model of that locus had been previously developed and shown to well reproduce Hi-C data^25^. The SBS is a model of chromatin describing the textbook picture where molecules, such as transcription factors, form DNA loops by bridging distal cognate binding sites^37^ (see also **Materials and Methods**). The SBS model has been shown to well describe Hi-C, GAM and FISH data across loci and cell types^20,24–27,39,45–47^. The genomic locations of the binding sites of the model of the *Sox9* locus were inferred from its Hi-C data^23^ by the PRISMR algorithm^25,27^, which finds the minimal set of binding sites (and cognate binders) best describing the input data from only polymer physics (**Materials and Methods**). Here, we considered the published model of the locus at 40kb resolution, with no additional refinements or improvements, and explored an ensemble of single-molecule 3D polymer structures derived by Molecular Dynamics simulations in the thermodynamics steady state of the system (**Figure 1a, Materials and Methods**).

**Figure 1.**
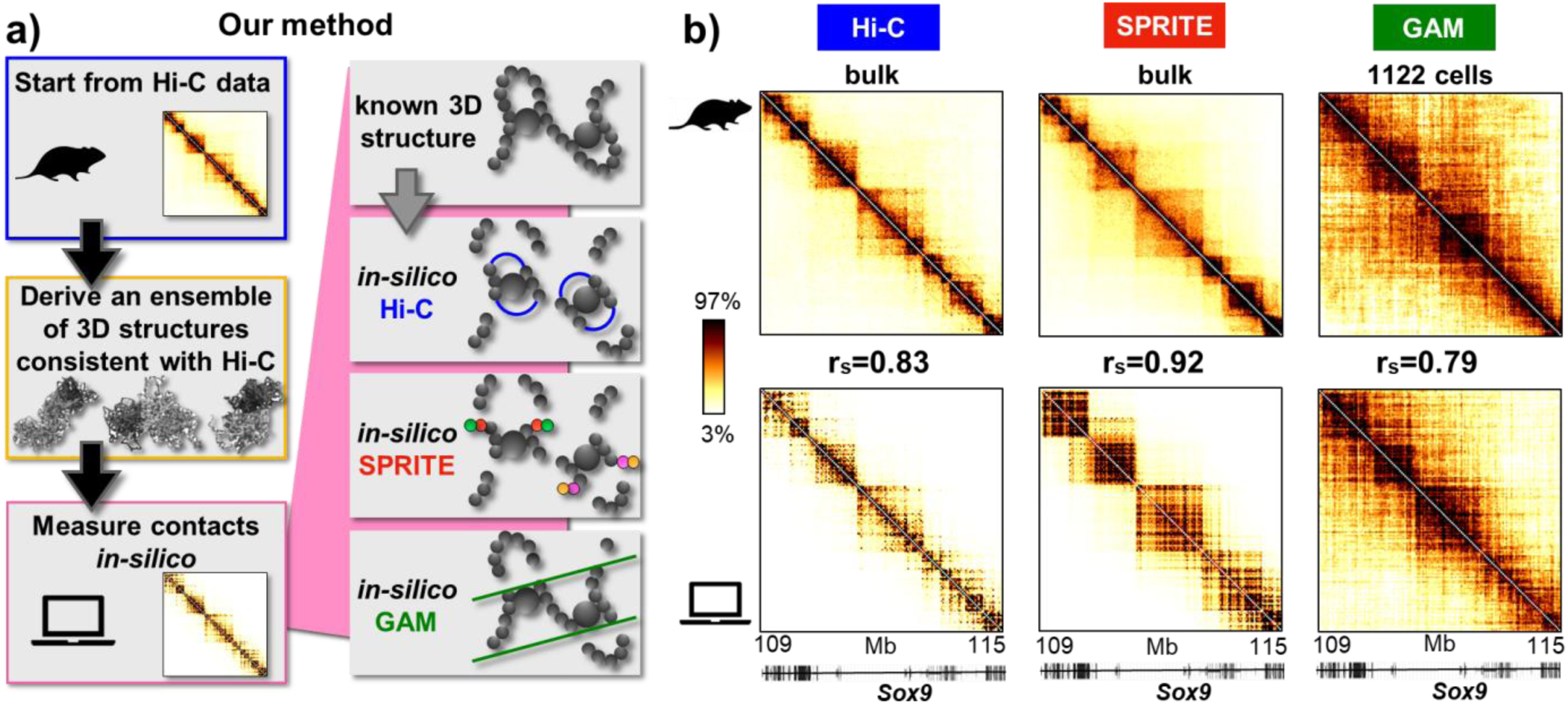
*In-silico* Hi-C, SPRITE and GAM average contact maps match experimental data. **a)** By use of the PRISMR procedure^27^, from Hi-C data the polymer model of the DNA locus of interest is inferred and, based on polymer-physics, an ensemble of its single-molecule 3D conformations, consistent with the input Hi-C data, derived. We implemented computationally the Hi-C, SPRITE and GAM methods on those 3D structures and measure *in-silico* the corresponding contact maps. **b)** Albeit inferred from Hi-C data only, the model 3D conformations return average contact maps (bottom) well matching the Hi-C and the independent SPRITE and GAM experimental data (top) in the case study of the *Sox9* locus (chr11:109Mb-115Mb, mm9) in mESC. Experimental Hi-C and SPRITE maps are bulk data^21,23^, while GAM data are from a new dataset constructed from 1122 F123 cells (**Materials and Methods**), and correspondingly the *in-silico* maps. The color scale represents the percentiles of each dataset. The Spearman correlation coefficients (r_s_) between model and experiment are reported in the middle, as Pearson or HiCRep correlations have similar values (**Supplementary Table 1a**).

Next, we computationally implemented the steps of the Hi-C, GAM and SPRITE methods on such ensemble of 3D structures to derive an *in-silico* proxy of their contact data (**Figure 1a**). In brief, in *in-silico* Hi-C, we fragmented in equal segments the two polymer chains representing the two *Sox9* alleles in each cell, ligated cross-linked fragments and counted ligation products to derive an *in-silico* analogue of Hi-C contact frequencies (**Materials and Methods**). The overall efficiency of the process is considered to be the product of the *in-silico* cross-linking, digestion, biotinylation, ligation and sequencing efficiencies. *In-silico* SPRITE was analogously implemented, in particular, by counting chain fragments tagged with the same barcode. Finally, *in-silico* GAM was performed by producing randomly oriented slices out of a sphere (representing the nucleus) where two single-molecule 3D structures (the two “alleles”) have been randomly positioned and by listing the polymer segments falling within each slice to derive the co-segregation matrix (see **Materials and Methods**). The overall efficiency here is the detection and sequencing efficiency of such segments, while the nuclear radius and the slice thickness are parameters set to match typical experimental values^20^ (**Materials and Methods**).

Such a procedure returns *in-silico* contact maps from the known 3D structures of the SBS model, providing a simplified, yet fully known benchmark to compare Hi-C, SPRITE and GAM in different contexts. In particular, we investigated how the overall efficiency and the number of pairs, N, of 3D single-molecule structures included in the analysis (below, N is named, for simplicity, the number of *in-silico* cells) affect the output of the three technologies.

### *In-silico* Hi-C, SPRITE and GAM reproduce experimental data

As our polymer model is inferred from Hi-C data^23^, we checked that the derived *in-silico* bulk Hi-C map, i.e., contact data averaged over the ensemble of 3D structures, reproduces bulk Hi-C experimental interaction frequencies in the *Sox9* locus in mESC^23^ (**Figure 1b**). We measured the correlation between the simulated and real Hi-C data and found that the Spearman (r_S_), Pearson (r) and HiCRep (scc)^48^ coefficients have all high values, respectively r_s_=0.83, r=0.83 and scc=0.80 (**Supplementary Table 1a**), as previously reported^25^. Analogous results were obtained in the comparison between *in-silico* and experimental Hi-C maps using the *HoxD* locus in mESC and the *Epha4* locus in CHLX-12 cells (**Supplementary Figure 1, Supplementary Figure 3a** and **Materials and Methods**).

Importantly, the *in-silico* SPRITE and GAM contact matrices derived from the same model 3D structures have also high correlations with the independent SPRITE and GAM experimental data (respectively r_s_=0.92 and r_s_=0.79, r=0.75 and r=0.80, and scc=0.57 and scc=0.40, **Figure 1b, Supplementary Table 1a**). In the comparison, we used published SPRITE bulk mESC data^21^ and a GAM dataset produced for the 4D Nucleome consortium^49^ (**Materials and Methods**) composed of 1122 nuclear profiles (slices) from F123 mESC cells, here compared with the output from precisely 1122 *in-silico* slices. Similar results are found for the mESC *HoxD* locus (**Supplementary Figure 1**). In particular, the visual differences and lower correlations between experimental and *in-silico* GAM contact matrices derived from Hi-C-based polymers may also raise the possibility that Hi-C and GAM experimental data may capture some different specific features of chromatin contacts, although they could just derive from experimental noise.

Overall, the agreement between model and experiments across independent datasets provides a validation of our polymer model, as the 3D structures inferred from Hi-C data only well reproduce independent GAM and SPRITE data too. It also shows that our *in-silico* approach has no major biases favouring Hi-C, SPRITE or GAM. That supports the view that our ensemble of 3D structures provides a good description of single-cell conformations of the loci and that the approach presented here can work as a simplified, yet useful reference system to compare the performance of the three technologies.

### Bulk Hi-C, SPRITE and GAM data all faithfully describe benchmark average distance matrices

Next, we investigated how well *in-silico* Hi-C, SPRITE and GAM data on the *Sox9* locus reflect the underlying spatial conformations of the polymers in the ensemble. Towards this aim, we computed the average distance matrix of the known 3D structures and compared it with *in-silico* Hi-C, SPRITE and GAM bulk data, i.e., the average over a large number of *in-silico* cells (**Figure 2**). We found that the three methods have high absolute Spearman correlation coefficients with the average distance matrix (r_s_<-0.89, values are negative because large physical distances correspond to small contact frequencies), GAM having the highest, followed by SPRITE and Hi-C (Pearson and HiCRep correlations give analogous results, **Supplementary Table 1b**).

**Figure 2.**
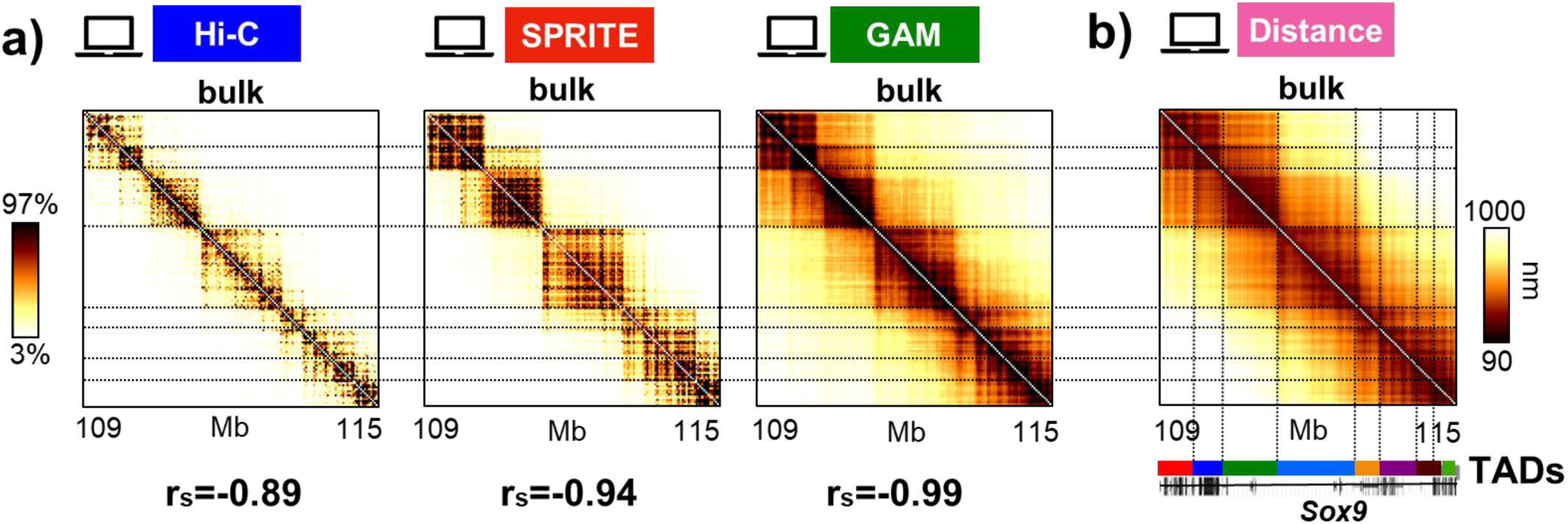
Bulk Hi-C, SPRITE and GAM data are all overall faithful to the average 3D distances. **a)** The *in-silico* bulk Hi-C, SPRITE and GAM maps of the *Sox9* locus, although corresponding to different measures of DNA physical proximity, return similar contact patterns. The color scale represents the percentiles of each contact map. The Spearman correlation coefficients are reported, on the bottom, between each contact map and the average 3D distance matrix in panel b) of the known single-molecule 3D conformations of the locus model (Pearson and HiCRep return similarly high correlations, **Supplementary Table 1b**). **b)** The average 3D distance map derived from the ensemble of *in-silico* model single-molecule 3D conformations is shown. Its high correlation values with each of three contact maps of panel a) (**Supplementary Table 1b**) illustrates that the three technologies faithfully capture the average pairwise distances of the system. We find that the TADs of the locus^23^ (different colors of the bar in the bottom) correspond to the domain-like patterns of the average distance matrix. They are well identified by the *in-silico* Hi-C, SPRITE and GAM bulk maps, as highlighted by the drawn vertical and horizontal lines. In particular, GAM captures well the longer-range inter-TADs contacts.

Interestingly, the patterns visible in the *in-silico* Hi-C, SPRITE and GAM bulk data are similar to each other, albeit GAM better highlights longer-range contacts between TADs (**Figure 2a**). In particular, the three *in-silico* methods all identify TADs that correspond to those previously found by experimental Hi-C^23^ (**Figure 2b**, different colours in the bottom bar). This is consistent with previous genome-wide analyses showing that the location of TAD boundaries in mESC is highly correlated in Hi-C, SPRITE and GAM^20,21^. Additionally, the contact patterns in Hi-C, SPRITE and GAM all reflect the underlying domain-like patterns of the average 3D distance matrix in the model locus, representing the known conformations of the ensemble of single-molecules inferred from Hi-C (**Figure 2b**). Again, analogous results are found for the *HoxD* and *Epha4* loci (**Supplementary Figures 2a,b** and **Supplementary Figures 3b,c**).

Taken together, our results support the view that bulk data from Hi-C, SPRITE and GAM are faithful to the overall spatial structure of the underlying 3D conformations, providing comparable information on the average distance map of the considered *Sox9, HoxD* and *Epha4* locus models.

### Stochasticity of single-cell data reflects intrinsic variability of single-molecule 3D conformations

Whereas bulk interaction matrices are comparatively similar across replicate experiments, single-cell Hi-C data exhibit a strong variability^50–54^. Here, we explore how single-cell variability reflects limited detection efficiency and, importantly, the inherent differences across single-molecule conformations of chromatin, whereby even single-cell experiments with 100% efficiency can return different contact maps (**Figure 3**).

**Figure 3.**
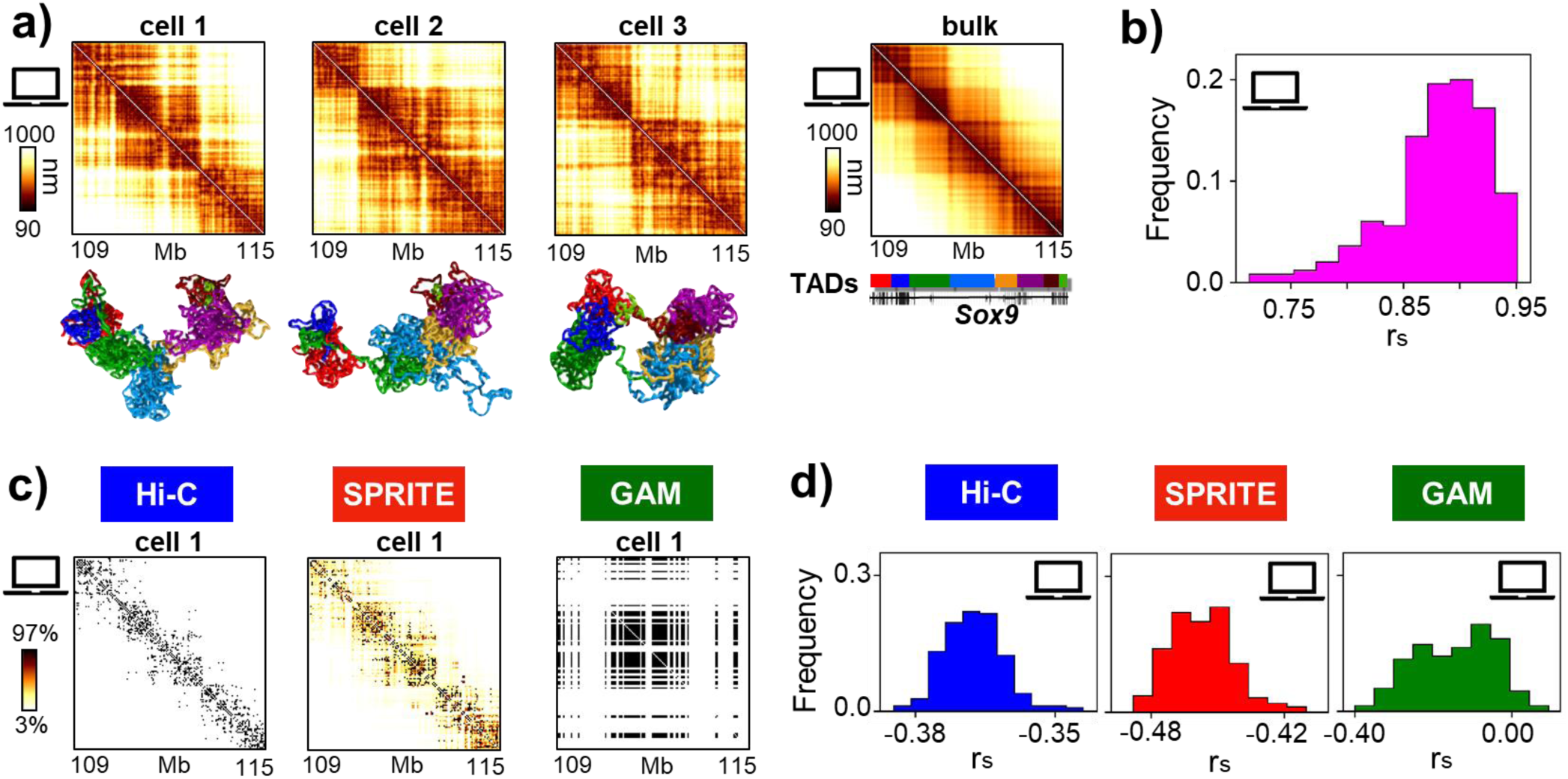
Stochasticity of single-cell contact maps reflects the intrinsic variability of single-molecule 3D conformations. **a)** The variability of single-molecule conformations of the model of the *Sox9* locus is represented here by three examples (bottom, the color scheme reflects the colors of the TADs of the locus^23^, shown in the color bar). Their corresponding *in-silico* single-cell distance maps (on top) can be different from the average distance matrix (left). For example, the Spearman correlations between the shown *in-silico* single-cell and the average distance maps are: r_S_=0.89 for cell 1; r_S_=0.82 for cell 2; r_S_=0.83 for cell 3. Replicate single-cell experiments can differ from each other just because they capture distinct underlying chromatin structures. **b)** The Spearman correlation between *in-silico* single-cell distance maps and the average distance map has a broad distribution (mean value r_S_ = 0.88 and median r_S_ = 0.89, **Materials and Methods**). Mean Pearson and HiCRep correlations are analogous and reported in **Supplementary Table 1c.** **c)** The *in-silico* single-cell Hi-C, SPRITE and GAM contact maps corresponding to the first of the three *in-silico* cells of panel a) are shown (color scale indicates the percentiles of each map). Here, the *in-silico* efficiency is set to 1, so all contacts are captured in Hi-C or SPRITE, and all segregated windows are detected in GAM. **d)** The distribution is shown of Spearman correlation coefficients between *in-silico* single-cell contact matrices at efficiency 1 and their corresponding *in-silico* single-cell distance matrices. *In-silico* single-cell data average correlations (**Supplementary Table 1d,e** and **Materials and Methods**) are overall significantly lower than correlations in bulk data (**Figure 2** and **Supplementary Table 1b**). Single-cell Hi-C and SPRITE perform comparatively better than single-cell GAM in reproducing the pattern of physical distances, as single-cell GAM derives from only a single slice cut out of a nucleus. Overall, single-cell contact matrices are less faithful to their corresponding distance map than bulk matrices (see **Figure 2**). Here we considered the ideal case where the efficiency is 1 to highlight the effects on contact data of the variability of single molecules, yet in real experiments the efficiency is different in the three technologies and well below 1 (see **Main Text** and **Figure 4**).

We discuss first the ideal case of *in-silico* experiments where the efficiency is set to 1.0. Single-molecule conformations vary widely across the ensemble of 3D structures (**Figure 3a, bottom**) and their corresponding *in-silico* single-cell distance matrices (**Figure 3a, top**) have broadly varying Spearman correlations with the average distance matrix, with a mean of r_s_=0.88 (**Figure 3b**, the values of the mean Pearson and HiCRep correlations are analogous, **Supplementary Table 1c**). Hence, there are broad structural differences between pairs of single molecules, as manifested in their corresponding single-cell measures. We found that the correlation of an *in-silico* single-cell Hi-C, SPRITE or GAM contact map (**Figure 3c**) with its corresponding single-cell distance matrix is much lower than in the case of bulk data discussed before: even in the case with efficiency 1, the average Spearman correlation is around r_s_=-0.37 and r_s_=-0.46 for respectively *in-silico* Hi-C and SPRITE (**Figure 3d**, and **Supplementary Table 1d**). For GAM the correlation is even lower (average r_s_=-0.15) and its distribution much broader, in the range −0.4 < r_s_ < 0. That is also a consequence of the different experimental procedure, because a single-cell *in-silico* Hi-C and SPRITE experiment returns the contacts measured over an entire *in-silico* nucleus, i.e., two independent polymer structures representing the alleles, whereas a single-cell *in-silico* GAM experiment probes the polymer content of only a single slice of an *in-silico* nucleus, i.e., a tiny fraction of the two polymers.

Summarizing, even in the ideal case of a 100% efficiency experiment, single-cell contact data are drastically less faithful to single-cell distance patterns than bulk data. In particular, single-cell *in-silico* Hi-C and SPRITE maps of the considered 6Mb locus have a correlation around r_s_=-0.4 with their corresponding single-cell distance matrices, whereas GAM has lower and much broader correlations because the experimental procedure samples a single slice rather than a single nucleus.

Contact data from single-cell experiments become further deteriorated, as expected, for lower values of the detection efficiency, and have worse correlations with the corresponding single-cell distance maps (**Supplementary Table 1e**). Consequently, the variability of replicates from *in-silico* single-cell experiments increases, i.e., the correlation between their contact maps decreases. For example, for an efficiency of 0.5, we found that the average correlation between *in-silico* single-cell replicates is around r_s_=0.2, 0.4 and 0.1 for respectively Hi-C, SPRITE and GAM maps. Importantly, the values of correlation found are consistent with those reported in real experimental studies: for example, the average Spearman correlation between different Hi-C maps of the *Sox9* locus from real single-cell experiments in CD4 T_H_1 cells with efficiency approximately around 0.025^50^ is r_s_=0.01, which is numerically equal to the Spearman correlation found between *in-silico* Hi-C maps at the same efficiency in our model of mESC (**Materials and Methods** and **Supplementary Figure 4a**). The impact of a limited efficiency on interactions maps is systematically investigated in the next section.

In brief, our results show that *in-silico* single-cell experiments are inherently broadly varying because they sample different single-molecule 3D structures and, additionally, their contact maps are less faithful to the corresponding single-cell distances. A limited detection efficiency further increases the fluctuations in contact maps to the point that replicates can have correlations well below 0.1 for realistic efficiencies. That appears to be reflected in, and consistent with, the stochastic nature of interactions between DNA sites observed in real single-cell experiments^20,50–53^.

### Threshold cell number required for replicate reproducibility differs in Hi-C, SPRITE and GAM

The quality of *in-silico* Hi-C, SPRITE and GAM contact maps improves when the number of *in-silico* cells, N, is increased in the computational experiments (**Figure 4**). Due to the intrinsic variability of single-molecule structures, the improvement with N occurs also in the ideal case of a 100% efficiency (**Figure 4a**). **Figure 4b** shows, for example, the effect of different N on the contact matrices of *in-silico* Hi-C, SPRITE and GAM, in the case where the analyses are run with efficiencies comparable to typical experimental values. We set the Hi-C efficiency to 0.05, taken as an upper limit of values reported in recent studies^50,51,53,55^; the same value is used as an estimate of the efficiency for SPRITE (M. Guttman, personal communication). Since the experimental efficiency in GAM is roughly one order of magnitude larger than Hi-C and SPRITE^20^, in the shown example we used an *in-silico* GAM efficiency of 0.5, a value which is close to the 0.6 efficiency estimated for the mESC dataset used in this work (**Materials and Methods**).

**Figure 4.**
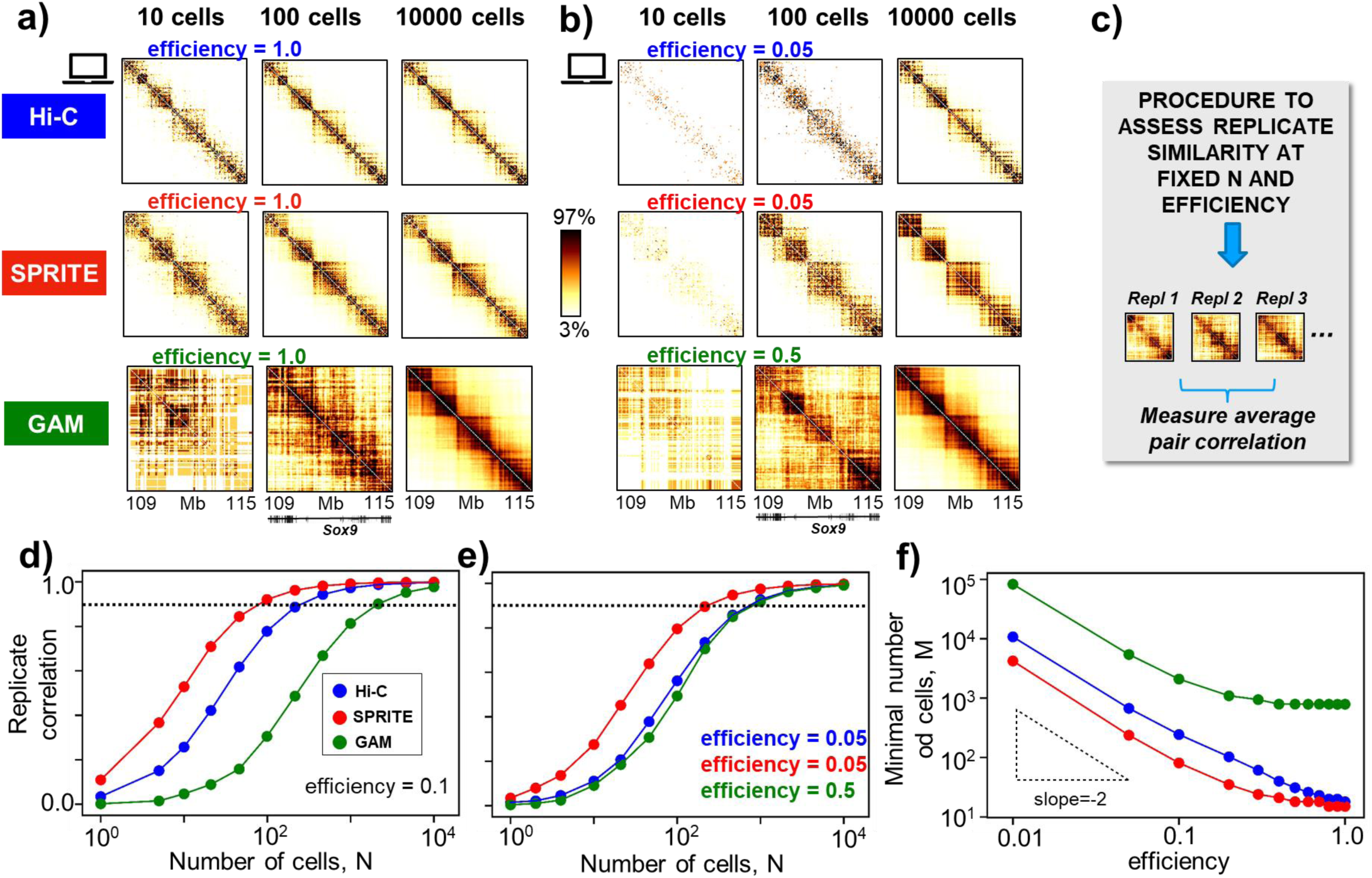
Threshold cell number required for replicate reproducibility differs in Hi-C, SPRITE and GAM. **a)** The *in*-*silico* Hi-C, SPRITE and GAM contact maps of the *Sox9* locus depend on the number of *in*-*silico* cells, N, considered in the experiment (here is shown the case with efficiency equal to 1), albeit in the bulk limit (large N), the effects of cell-to-cell variability are averaged out for all the three technologies. Color scale indicates the percentiles of each map. **b)** Results analogous to those in panel a) are shown in the case where efficiencies similar to those found in real experiments are considered: here, for Hi-C and SPRITE the *in-silico* efficiency is set equal to 0.05, and for GAM equal to 0.5 (see **Main Text**). **c)** To assess the similarity between *in-silico* replicate contact maps (i.e. maps obtained with same number of cells, N, and efficiency), we measured the average Pearson correlation between them. The minimal number of cells, M, to have a reproducible output map is defined as the value of N where the average Pearson correlation between replicates crosses the threshold r_t_=0.9 (**Main Text** and **Materials and Methods**). **d)** The Pearson correlation is shown between replicate experiments as a function of N for Hi-C, SPRITE and GAM at a given efficiency (0.1). The dashed line is the threshold correlation value r_t_=0.9. Analogous results are found when the Spearman or HiCRep correlations are considered (**Supplementary Figure 5**). **e)** Results analogous to those in panel d) are shown in the case of efficiencies similar to those in real experiments, as discussed in panel b). As GAM has a higher efficiency, its corresponding behavior with N becomes closer to those of Hi-C and SPRITE. For example, here, for the reported realistic efficiencies of 0.05 for Hi-C and SPRITE, and 0.5 for GAM, M is respectively approximately 650, 250 and 800. **f)** The value of M is shown for Hi-C, SPRITE and GAM as a function of the efficiency. M increases as the efficiency is reduced and grows approximately as an inverse squared power law at small efficiencies. The obtained values of M are consistent with the Central Limit Theorem (**Materials and Methods**). For a given efficiency, M is the smallest in SPRITE, a factor of two higher in Hi-C and one order of magnitude higher in GAM.

The patterns of the contact matrices become sharper and stabilise when N becomes large enough, as also observed in experimental investigations^50,56^. Importantly, overall the large N matrices do not depend on the considered efficiency value and the average over a large number of cells compensates, in general, for reduced efficiencies (see **Figure 4a** and **4b** and **Materials and Methods**). However, our data show that the threshold value of N to reach saturation in the data strongly depends on the efficiency level and is different in different technologies, as we now illustrate.

We aimed to identify the minimal number of cells that, at a given efficiency, is required for replicate experiments to return similar outputs, i.e., to approach the bulk limit, in the *in-silico* implementation of Hi-C, SPRITE and GAM. To measure the similarity of pairs of identical experiments in the *Sox9* locus (each having a given N and efficiency, **Figure 4c**), we computed the average Pearson correlation between contact maps (**Figures 4d,e**; the use of Spearman or HiCRep correlations returned analogous results **Supplementary Figure 5**). The correlation grows when N is increased and it plateaus to 1 in the large N limit (**Figures 4d,e**), independently of the efficiency of the *in-silico* experiment. For each given efficiency, we heuristically define the minimal number of cells, M, required for having statistically reproducible results across replicates, as the value of N where the correlation grows larger than a given threshold, r_t_=0.9 (**Materials and Methods**). Importantly, we found that M is significantly different in the different technologies: for example, if the efficiency is 0.1, we found that M is 200, 100 and 2000 for respectively Hi-C, SPRITE and GAM (**Figure 4d**). **Figure 4e** shows the correlation between replicates at varying N obtained for efficiencies close to those reported in real Hi-C, SPRITE and GAM experiments, i.e., as specified above, 0.05 for Hi-C and SPRITE and 0.5 for GAM: in those cases, M is approximately 650, 250 and 800 respectively. Similar behaviours were also found for the *HoxD* and *Epha4* loci (**Supplementary Figures 2c** and **3d**). In general, for a given efficiency, we find that *in-silico* SPRITE plateaus earlier than Hi-C, while GAM typically requires a number of cells one order of magnitude larger. However, GAM experiments have currently efficiencies one order of magnitude higher than Hi-C or SPRITE, hence the number of cells required for saturation become similar in the three methods.

Next, we checked how the *in-silico* estimates of M compares against available systematic experimental investigations. First, we verified that the correlation values between *in-silico* replicates are comparable to those found in experiments. We considered the 60 different single-cell Hi-C maps produced in CD4 T_H_1 cells^50^ with efficiency around 0.025 and compared their average map with the corresponding bulk Hi-C data for the *Sox9* locus: the correlation is r_s_=0.33, which is not far from r_s_=0.27 found between *in-silico* Hi-C maps in the analogous conditions in mESC (**Supplementary Figure 4b** and **Materials and Methods**). Second, to verify that our estimates of M are consistent with available experimental results we considered the data from a recent Low-C experiment on mESC^56^ where, in the case of a 10Mb wide locus, a sample of 1000 cells was shown to be large enough to produce contact maps highly similar to the bulk one (Pearson correlation r=0.95). That estimate of the minimal number of cells needed to approach the bulk limit is consistent with the above reported value of M=650 for *in-silico* Hi-C for an efficiency equal to 0.05. Taken together, these examples show that the *in-silico* estimates of M are informative of real experiments.

Finally, we systematically investigated how the quality of the *in-silico* data is affected by the efficiency of the experiment (**Supplementary Figure 6**). In particular, the number of cells required for saturation, M, strongly depends on the efficiency (**Figure 4f**): M diverges approximately as an inverse squared power law as the efficiency becomes small. In other words, halving the efficiency requires to quadruple the cell number to achieve the same quality levels. In general, we find that M for SPRITE is two times smaller than the corresponding value for Hi-C and one order of magnitude smaller than GAM. Additionally, our investigation shows that even in the ideal case of an efficiency equal to 1, single-cell replicates have below threshold correlations, as M is larger than 10 even for SPRITE due to the intrinsic variability of single-molecule 3D structures, as reported above. That rationalises of the broad variability observed in single-cell Hi-C experiments.

We stress that the above definition of M is heuristic, albeit easy to visualise. It is, however, fully consistent with a definition grounded on the Central Limit Theorem (CLT). Consider the average value, μ, and the standard deviation, σ, of a given entry of a contact map in an experiment with N cells at a given efficiency. CLT imposes that the noise-to-signal squared ratio, σ^2^/μ^2^ scales as 1/N. Accordingly, from the CLT, the minimal number of cells, L, required to make the noise-to-signal squared ratio smaller than a given threshold, δ, is L=Aδ^-2^σ^2^/μ^2^, where A is a constant (**Materials and Methods**). We checked that M and the average value of L are linearly proportional to each other, i.e., M is inversely proportional to the squared signal-to-noise ratio averaged over all the entries of a single-cell contact map, *ρ* (**Materials and Methods** and **Supplementary Figure 7**). Hence, albeit heuristic, the above intuitive definition of M is grounded on the CLT. With analogous statistical arguments (**Materials and Methods**) the approximate inverse squared power law relation between M and the efficiency (**Figure 4f**) can be explained.

The sets of *in-silico* single-molecule 3D structures employed in all our analyses were produced using polymer models inferred from Hi-C data^25–27^. However, for the *Sox9* locus we tested that our results remain overall unchanged also when the polymer 3D structures are constructed from the SBS polymer model of the locus inferred from GAM data rather than Hi-C^45^ (**Supplementary Figure 8** and **Materials and Methods**). Additionally, to assess the general validity of our analyses, we applied the *in-silico* approach to 3D conformations of a toy block-copolymer, unrelated to real chromatin loci, finding similar results (**Supplementary Figure 9** and **Materials and Methods**).

### SLICE single-cell interaction probability maps

Next, we investigated the performance of the GAM data analysis tool SLICE (**Supplementary Figure 10**). SLICE is a statistical method to identify non-random co-segregation events (i.e., specific interactions) from GAM co-segregation data. In particular, the output of SLICE is the single-cell interaction probability (Pi) of pairs, and multiplets, of DNA sites^20^. Importantly, we found that SLICE bulk interaction probabilities are faithful to the known average distance matrix (r=-0.95, r_s_=-1.00, scc=-0.99, **Supplementary Figures 10a,b**). The SLICE matrices behave with N and with the efficiency as found for GAM contact maps (**Figure 4**). However, as by definition SLICE specifically detects significant interactions, we found that the average number of *in-silico* cells, M, needed to return statistically reproducible results across replicates is approximately half than the one required for GAM in the same conditions (**Supplementary Figures 10c-e**). For a realistic efficiency of 0.5, for example, we found that M=400 for SLICE, whereas M=800 for GAM. In that respect, SLICE can be employed as an useful tool in applications of GAM, especially where the number of available cells is small, such as in the analysis of sample tissues or biopsies.

Summarizing, our findings illustrate how the level of variability of *in-silico* contact matrices is affected by the number of single cells, N, and by the experimental efficiency, and how different technologies perform in different situations. Overall, under same conditions, SPRITE turns out to be the more suitable method to extract pair-wise contact information from small cell samples as M is smaller than in Hi-C, and GAM requires a much larger number of cells for replicate robustness. Interestingly, however, at realistic efficiency values for Hi-C, SPRITE and GAM, the number of required cells become similar across the three methods, especially if SLICE is used in combination with GAM.

### Noise-to-signal levels vary with genomic distance differently in Hi-C, SPRITE and GAM

Finally, we investigated the noise-to-signal level of the entries of contact matrices and how it varies with the genomic separation, with the number of cells, N, and with the efficiency of *in-silico* experiments. For each entry of a contact map, as discussed before, the noise-to-signal ratio is defined as the ratio of the standard deviation, *σ*, to mean value, μ, across replicates from experiments in the same conditions. For a given N and a given efficiency, we observed that the average noise-to-signal ratio, ⟨*σ*/*μ*⟩, is strongly dependent on genomic distance (**Figure 5a**). In the *Sox9* locus, we found for both Hi-C and SPRITE that ⟨*σ*/*μ*⟩ grows more than one order of magnitude as the genomic separation increases from 50kb to 5Mb. In particular, there is a steep increase above the 1Mb scale. SPRITE has the lowest ⟨*σ*/*μ*⟩ ratio at genomic scales below the Mb, but interestingly GAM has an overall less varying noise-to-signal level. This is deriving from the GAM methodology that in a single slice samples DNA regions spanning the entire nucleus. Therefore, at larger genomic separations, GAM has almost a one order of magnitude lower noise-to-signal ratio than Hi-C and SPRITE.

**Figure 5.**
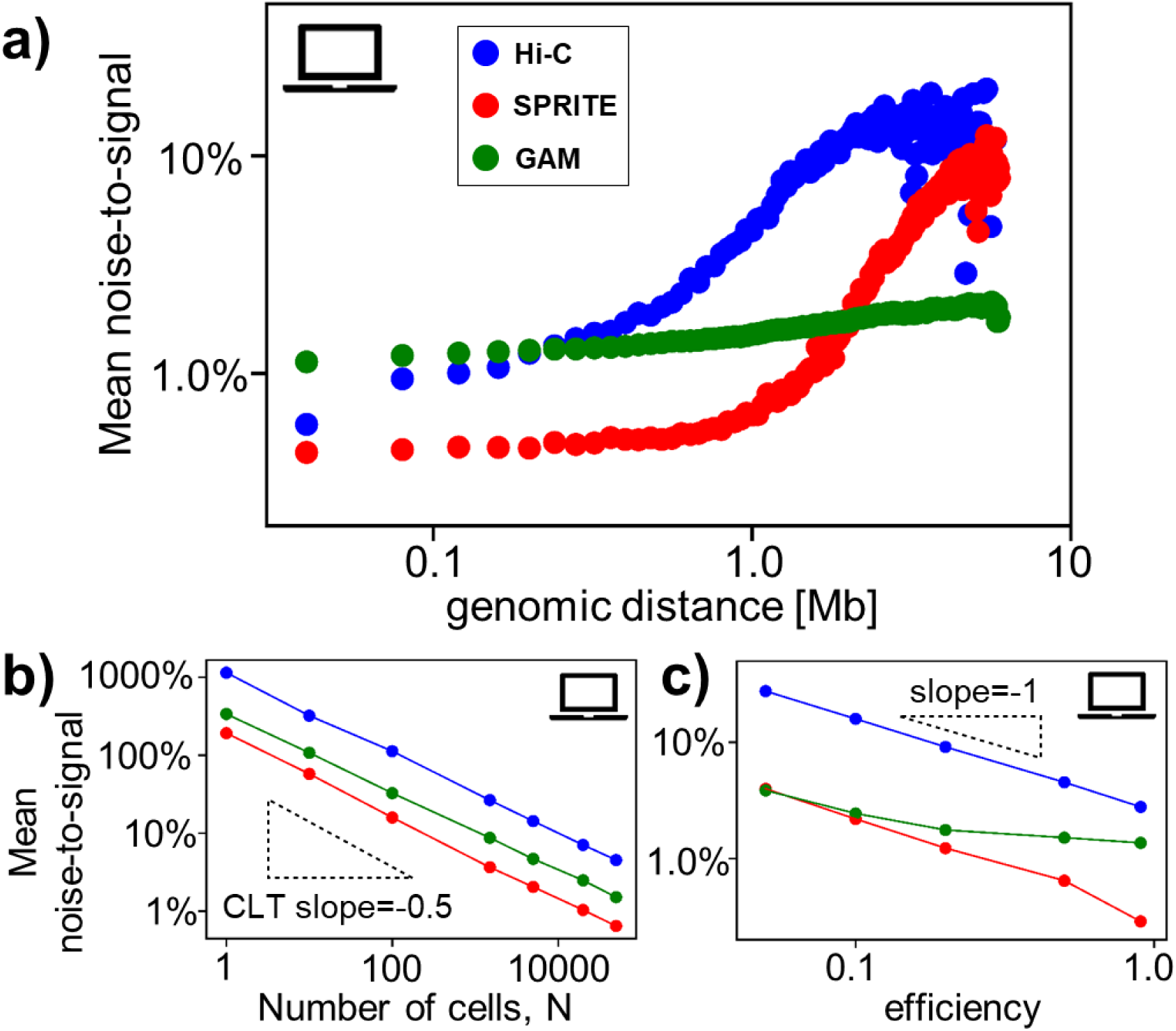
Noise-to-signal levels vary with genomic distance differently in Hi-C, SPRITE and GAM. **a)** The mean noise-to-signal ratio ⟨*σ*/*μ*⟩ of a contact map (see **Main Text** and **Materials and Methods**), for a given number of cells N and efficiency, depends on the considered genomic separation. For the *Sox9* locus, in Hi-C and SPRITE, the noise-to-signal ratio drastically grows above 1Mb, i.e. the scale of TADs, while GAM retains low noise-to-signal ratios across larger genomic separations (the case shown is for N=50000 and efficiency=0.5). **b)** For a given genomic distance and efficiency (the case shown is for 1Mb and efficiency=0.5), ⟨*σ*/*μ*⟩ decreases with N as an inverse square root, as expected from the Central Limit Theorem. **c)** Analogously, for a given genomic distance and N (the case shown is for 1Mb and N=50000), ⟨*σ*/*μ*⟩ increases, for small efficiencies, approximately as an inverse power law when the efficiency is reduced.

At a given genomic distance and efficiency, as expected, the noise-to-signal ratio decreases as the number of cells, N, is increased in our computational experiments (**Figure 5b**). Consistent with the Central Limit Theorem, it follows an inverse squared power law in N (i.e., N^-1/2^). A consequence of such a scaling behaviour is that single-cell (N=1) contact maps become highly noisy at large genomic separations. For example, at the 1Mb scale and for a detection efficiency of 0.5, the noise-to-signal ratio for N=1 is larger than 100% for all three methods, Hi-C having the largest fluctuations with ⟨*σ*/*μ*⟩>1000%. As expected, the noise-to-signal ratio is also strongly affected by the experimental efficiency (**Figure 5c**): in brief, we find that for a given genomic distance and for a given N, ⟨*σ*/*μ*⟩ decreases roughly as an inverse power law of the efficiency in our *in-silico* study.

In summary, we find that GAM is the method comparatively less noise affected at larger genomic distances, while SPRITE has the best performance below the Mb scale.

## DISCUSSION

Hi-C, SPRITE and GAM are powerful technologies to generate genomic contact maps, which return different measures of physical proximity between DNA sites, each affected by specific biases and limitations. As we lack an experimental benchmark to compare their performance, in this study we ran a computational investigation where we implemented *in-silico* the three methods on ensembles of known single-molecule 3D structures to compare them within a simplified, yet fully controlled framework. In particular, we focused, as a case study, on a 6Mb wide region around the *Sox9* gene in mESC. To check the robustness of our main conclusions, we also investigated a region around the *HoxD* genes in mESC and around the *Epha4* gene in CHLX-12 cells, as well as a toy block-copolymer model, finding analogous results.

The agreement between independent Hi-C, SPRITE and GAM bulk data and *in-silico* maps in all the studied loci supports the use of our single-molecule 3D structures and the *in-silico* method as a proxy for real experiments, providing a tool to compare those technologies in a fair way. Importantly, in all the studied cases, we find that *in-silico* Hi-C, SPRITE and GAM bulk contact data, as well as SLICE interaction probabilities, all faithfully represent the known spatial conformations of the model polymers. That also suggests that our approach has no biases in favour of any of the three technologies. In particular, we observed that the different methods identify very similar pairwise contact patterns, such as TADs and sub-TADs, which are found to correspond to the known underlying structure of the 3D conformations of the polymer ensemble.

We analysed how the entries of the contact maps of Hi-C, SPRITE and GAM are differently affected by some important parameters of the computational experiments, such as the number of *in-silico* cells, N, the detection efficiency and the genomic distance. We also quantified how the stochasticity of single-cell experiments is dependent on the intrinsic variability of chromatin 3D conformations. In summary, we showed that if N is below a threshold value M, replicate experiments can return broadly different outcomes. The value of M, consistent with arguments based on the Central Limit-Theorem, increases as the efficiency of the experiment decreases. For equal conditions, M is different in different technologies: SPRITE is the method having the lowest M and so better performing on samples with a small number of cells; GAM has the highest value, but in combination with SLICE M is drastically reduced. In real applications, it is important to take into account that the efficiency is different across the three methods, and the corresponding values of M can become similar. For example, the experimental estimations of the efficiency of Hi-C is around 0.05^50^, SPRITE is around 0.05 too (M. Guttmann personal communication) and for GAM around 0.5^20^: in those considitions we find that M is around 650, 250 and 800 for respectively Hi-C, SPRITE and GAM; additionally, when GAM is combined with SLICE, M becomes approximately 400. Reassuringly, we found that the use of large N can generally compensate for a limited efficiency and in the bulk limit the different technologies are all overall faithful to the benchmark model 3D structures. Finally, we analysed the noise-to-signal ratio in contact maps as a function of the genomic distance and found that GAM is less noise sensitive at large genomic separations, while Hi-C and SPRITE at lower distances, under the same conditions.

More generally, the consistent behaviour of our computational analyses across all the investigated cases (models of real loci as well as toy models) supports the view that their conclusions have a broad validity and can help guiding the design of novel Hi-C, SPRITE and GAM experiments in different contexts and applications.

## ACKNOWLEDGEMENTS

M.N. acknowledges support from CINECA ISCRA ID HP10CYFPS5 and HP10CRTY8P, Einstein BIH Fellowship Award (EVF-BIH-2016-282), Regione Campania SATIN Project 2018-2020, and computer resources from INFN, CINECA, ENEA CRESCO/ENEAGRID^57^ and *Scope*/*ReCAS* at the University of Naples. A.P., R.K., A.K., I.I.A thank the Helmholtz Association (Germany) for support. M.N. and A.P. acknowledge support from the National Institutes of Health Common Fund 4D Nucleome Program grant U54DK107977 and the EU H2020 Marie Curie ITN n.813282. I.I.A. acknowledges support from the Federation of European Biochemical Societies (FEBS Long-Term Fellowship).

## AUTHOR CONTRIBUTIONS

M.N. designed the project with inputs from A.P.. M.N., L.F. and F.M. developed the modelling. L.F. and F.M. ran computer simulations and performed data analyses with help from S.B., A.M.C., A.E. and M.C.. R.K., A.K. and I.I.A. produced and normalized the GAM dataset. M.N., L.F., F.M. and A.P. wrote the manuscript, with inputs from all the authors.

## SUPPLEMENTARY FIGURES

**Supplementary Figure 1.**
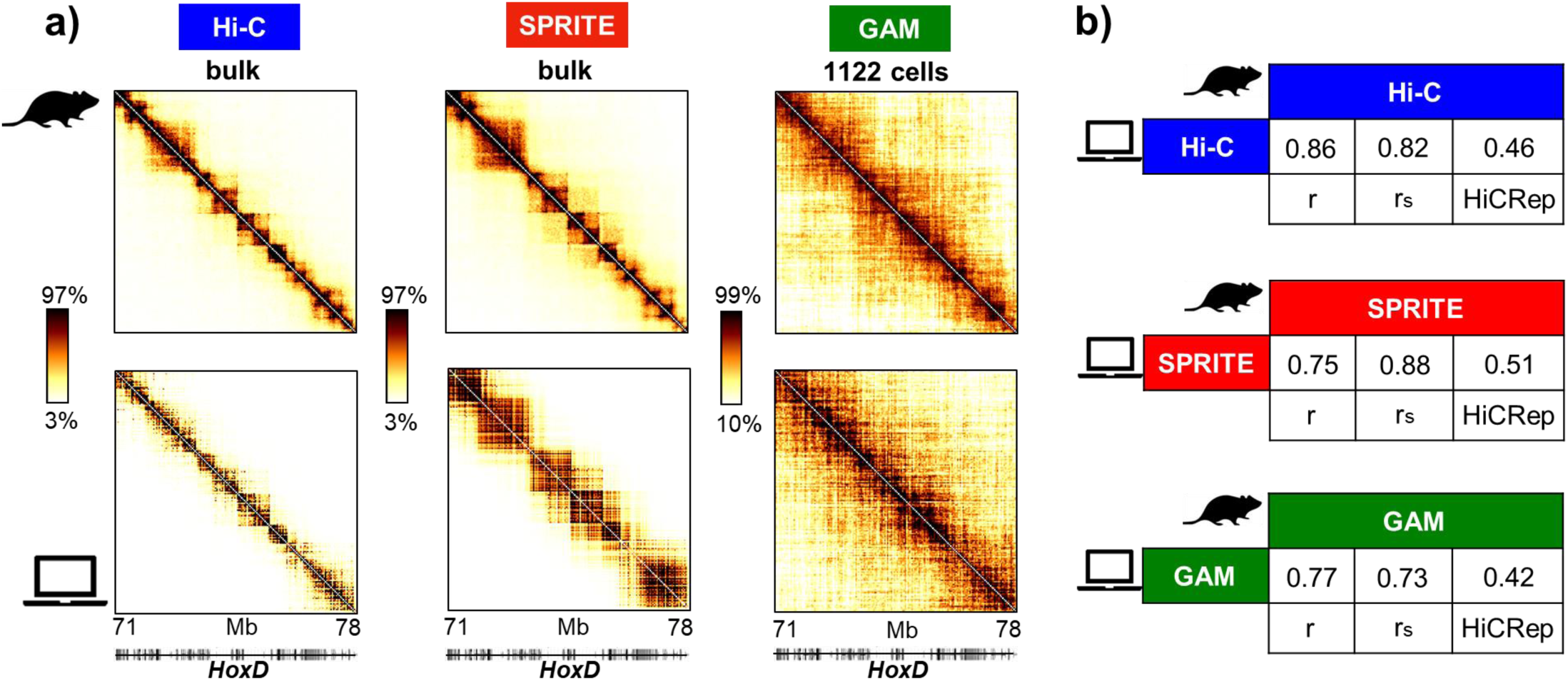
*In-silico* average contact maps of the *HoxD* locus compare well with Hi-C, SPRITE and GAM experimental data. a) In the case of the murine *HoxD* locus (chr2:71Mb-78Mb, mm9) in mESC, our 3D conformations return *in-silico* contact maps (bottom) that match well with Hi-C^23^, SPRITE^21^ and GAM experimental data from mESC (top). GAM data are from the new dataset of 1122 nuclear slices (**Materials and Methods**). Correspondingly, *in-silico* Hi-C and SPRITE are bulk data, while *in-silico* GAM data are from 1122 *in-silico* cells (see **Materials and Methods**). Color scale indicates the percentiles of the maps. b) Pearson, Spearman and HiCRep correlations between the *in silico* and experimental maps of panel a), for the three technologies.

**Supplementary Figure 2.**
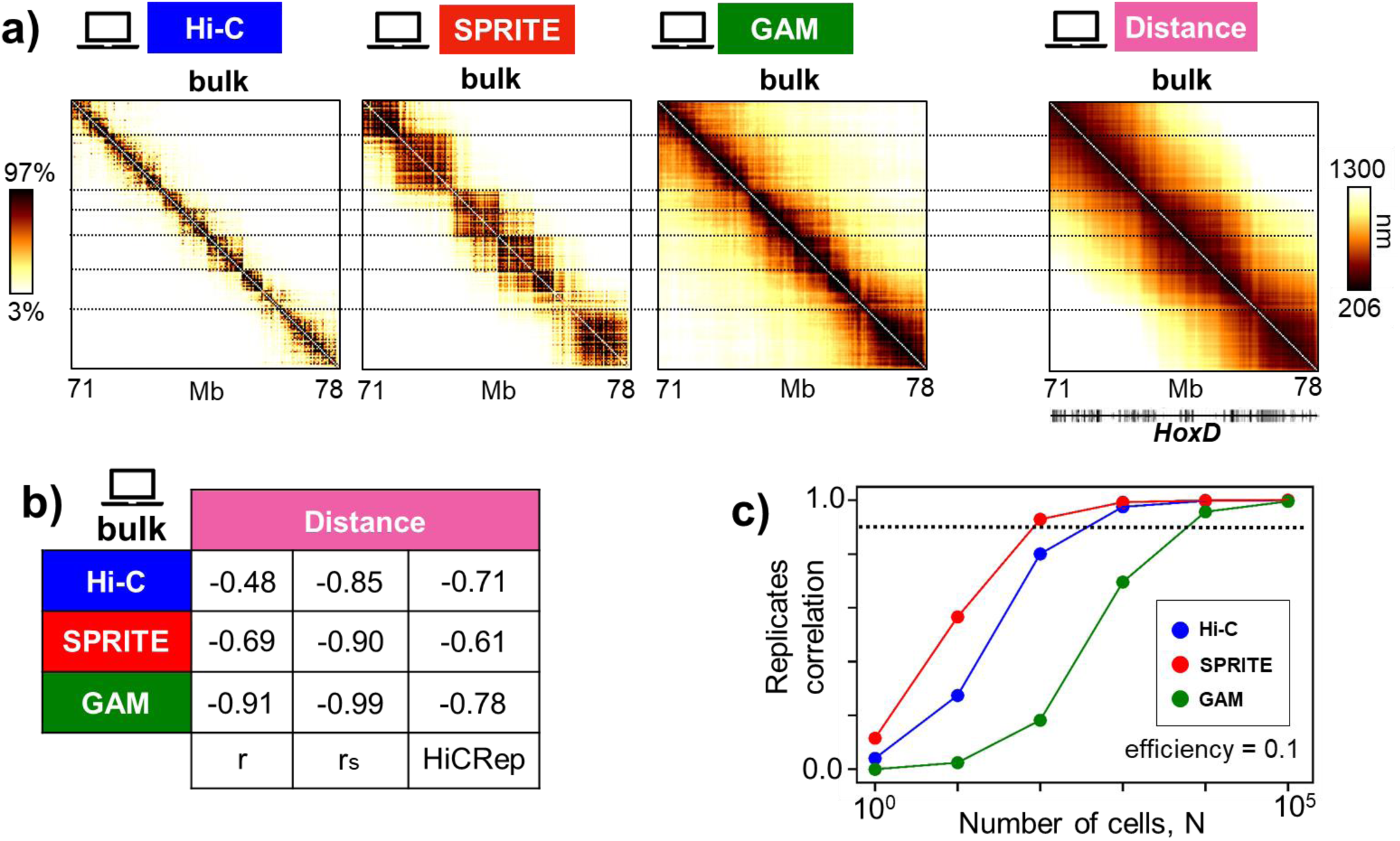
*In-silico* bulk contact maps overall reproduce the average distance map in the *HoxD* locus. a) *In-silico* bulk Hi-C, SPRITE and GAM maps of the *HoxD* locus return contact patterns that are compatible with the average distance pattern derived from the ensemble of single-molecule 3D conformations. The horizontal lines are drawn to mark the domain-like structure of the distance map. The color scale indicates the percentiles. b) Pearson, Spearman and HiCRep correlations are reported between the bulk contact maps and the average distance map. Correlations indicate a high degree of similarity, showing that the three technologies all faithfully capture the underlying conformations of the *HoxD* locus, analogously to the *Sox9* case study (**Figure 2** and **Supplementary Table 1b**). c) The Pearson correlation is shown between replicates against the number of *in-silico* cells for the *HoxD* locus at efficiency 0.1. The dashed line is the threshold correlation value r_t_=0.9. The resulting trend is similar to the *Sox9* locus case in **Figure 4d.** Also, the M values found for Hi-C, SPRITE and GAM are analogous to the *Sox9* ones in **Figure 4d**.

**Supplementary Figure 3.**
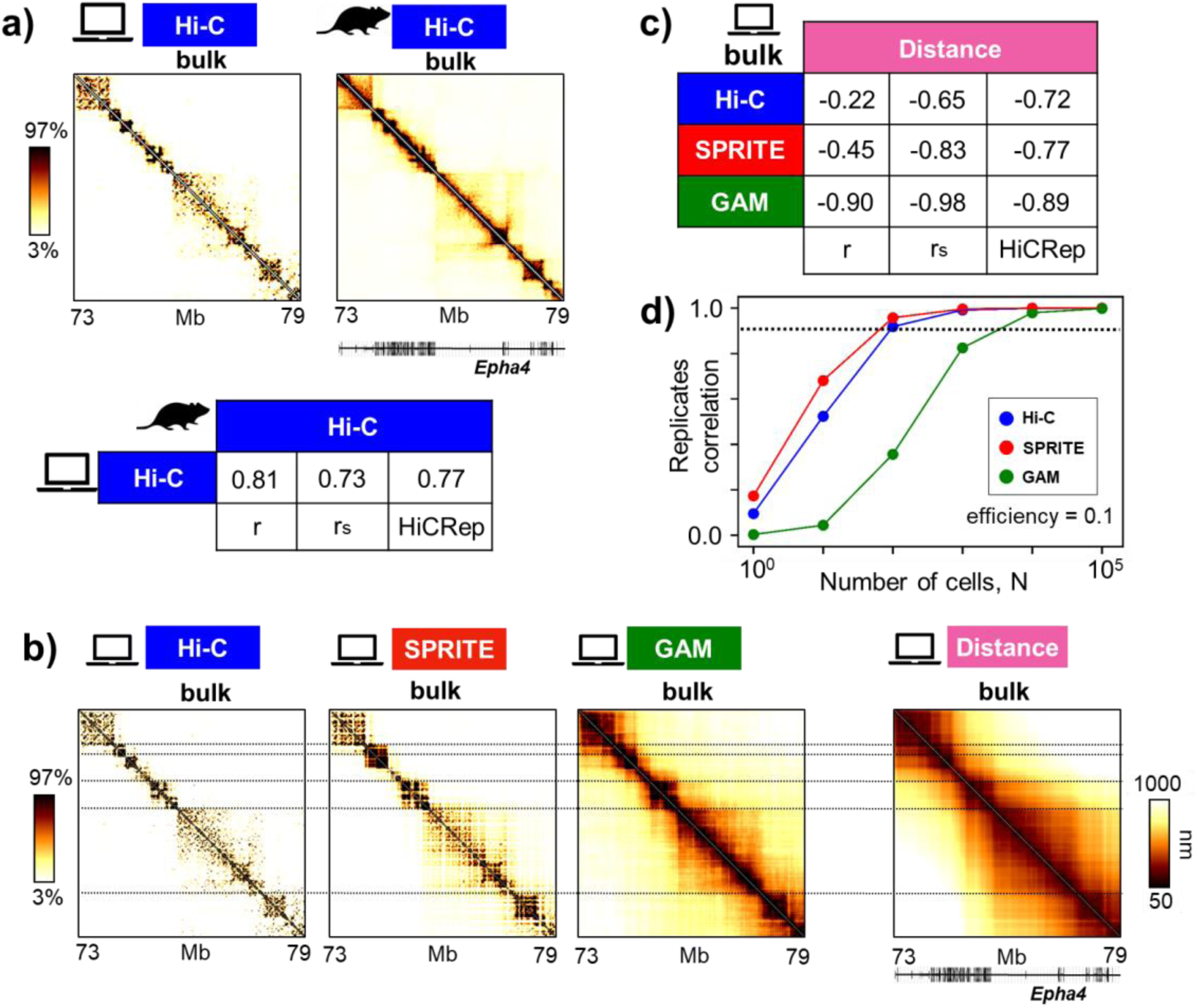
*In-silico* contact maps from 3D conformations of the *Epha4* locus. a) In the model of the *Epha4* locus in mouse CHLX-12 cells the *in-silico* bulk Hi-C map (left) is compared to the experimental map^15^ (right) (color scale indicates the percentiles of the maps). In the table on the bottom, Pearson, Spearman and HiCRep correlations are reported, indicating good similarity between the two matrices. b) The *in-silico* bulk contact maps are compared with the average distance pattern obtained from the ensemble of 3D conformations of the model of the locus. The horizontal lines are drawn to mark the domain-like structure of the distance map. c) Pearson, Spearman and HiCRep correlations are reported between each bulk contact map and the average distance map, indicating overall a good degree of similarity for each of the technologies. d) The Pearson correlation is shown between replicates against the number of *in-silico* cells for the *Epha4* locus, at efficiency 0.1. The dashed line is the threshold correlation value r_t_=0.9. The values of M are compatible with the ones we find in the *Sox9* and *HoxD* loci (**Figure 4d** and **Supplementary Figure 2c**), for all three technologies.

**Supplementary Figure 4.**
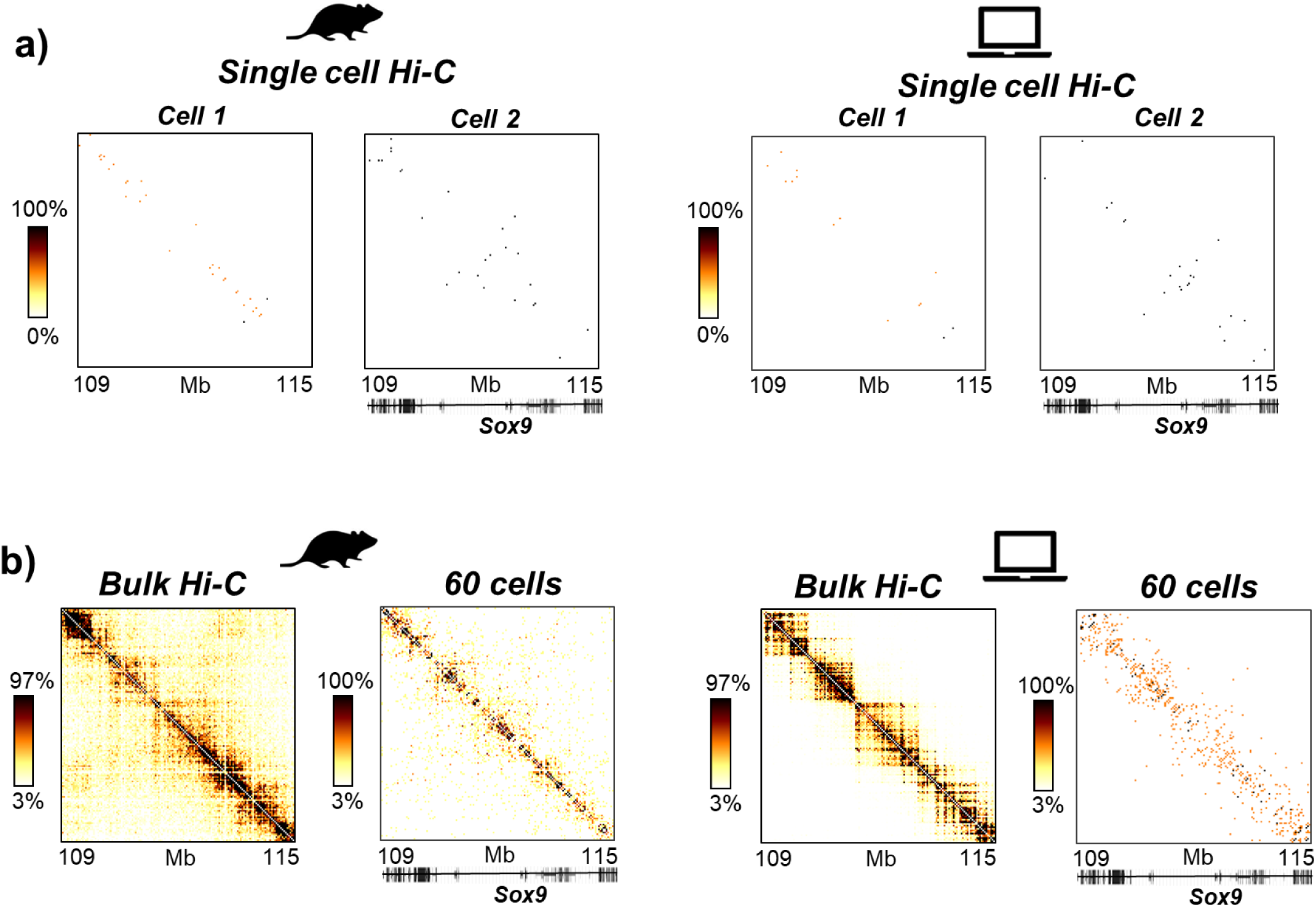
Experimental and *in-silico* single-cell Hi-C data. a) Left panel. Two examples of experimental single-cell Hi-C contact maps^50^, for the *Sox9* locus in the mouse CD4 T_H_1 cells. The mean Spearman correlation between all the available pairs of such single-cell *Sox9* maps^50^ is rS=0.01 (see also **Materials and Methods**). Right panel. Two examples of the *in-silico* single-cell Hi-C maps for the *Sox9* locus in mESC. Mean Spearman correlation is r_S_=0.01, consistent to the experimental result (**Materials and Methods**). The efficiency is set to 0.025^50^. Color scale indicates the percentiles of the maps. b) Left panel. In the same genomic region in CD4 T_H_1 cells, the average experimental map resulting from 60 available single-cell contact data is compared against the bulk Hi-C map^50^: their Spearman correlation is r_S_=0.33 (see **Materials and Methods**). Right panel. A similar calculation from the *in-silico* Hi-C maps in mESC returns a Spearman correlation between the bulk and the 60-cell map of r_s_=0.27, close to the experimental value (**Materials and Methods**). Color scale indicates the percentiles of the maps.

**Supplementary Figure 5.**
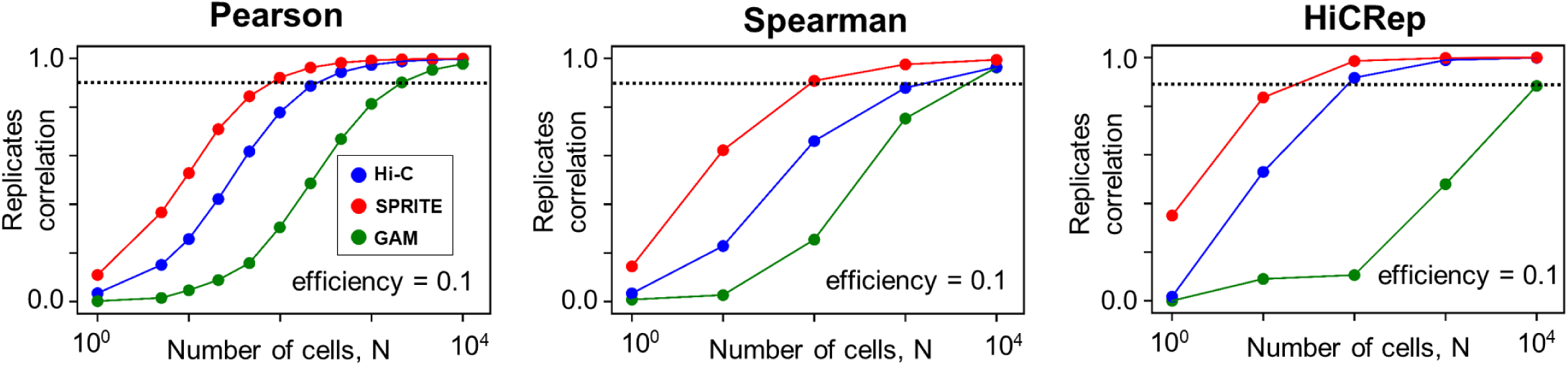
Pearson, Spearman and HiCRep correlations between replicates in relation to the number of cells considered in the *in-silico* experiments. The Pearson, Spearman and HiCRep correlations between replicate *in-silico* contact maps are shown for Hi-C, SPRITE and GAM at efficiency 0.1, in the case of the model of the *Sox9* locus. Dashed lines in each plot indicates the considered 0.9 threshold value (see **Figures 4c,d,e**).

**Supplementary Figure 6.**
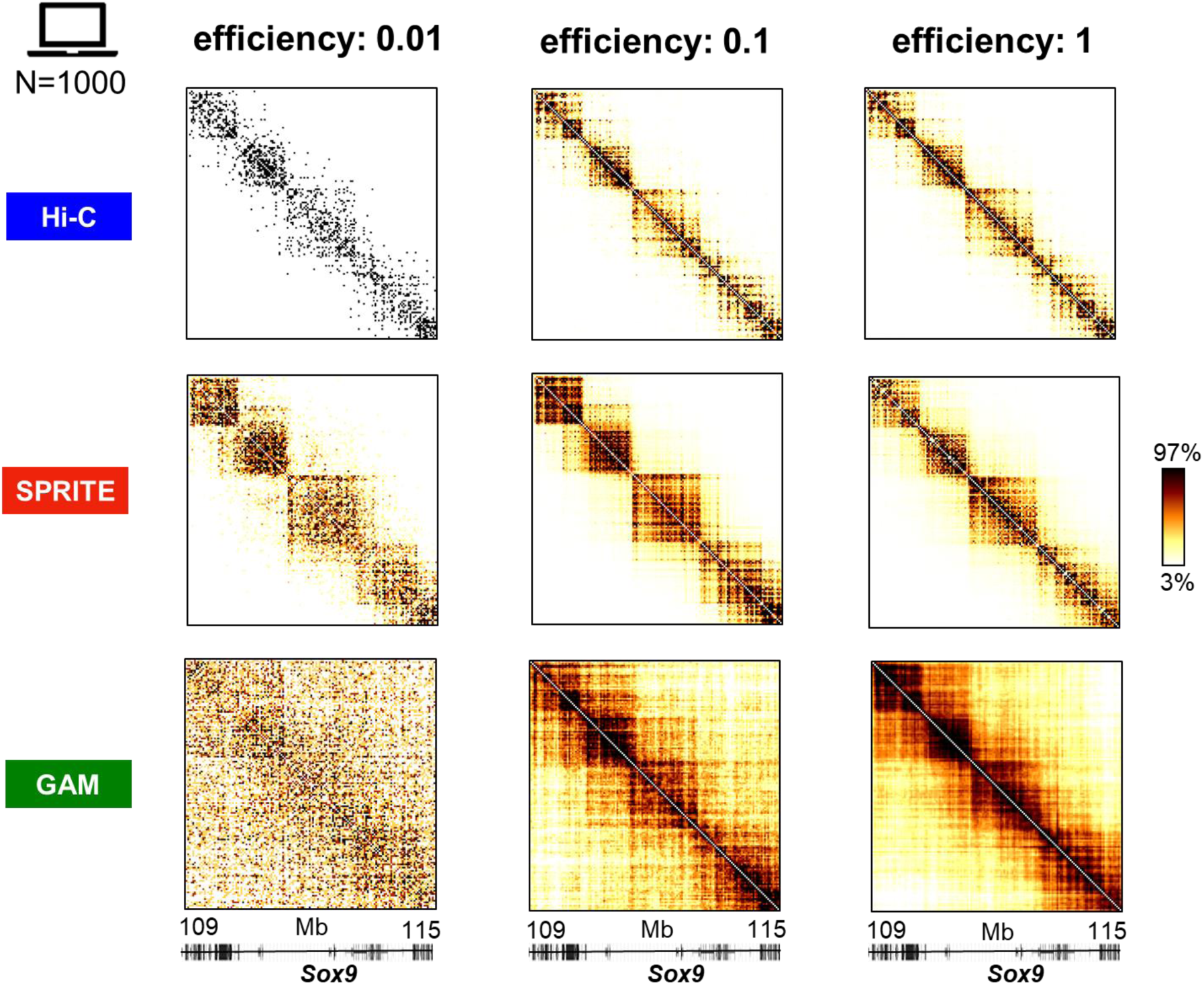
Impact of the detection efficiency on *in-silico* contact maps. Hi-C, SPRITE and GAM *in-silico* contact maps are shown for three different efficiencies (0.01, 0.1 and 1.0) for a fixed N=1000 *in-silico* cells, in the *Sox9* locus case study. Low efficiencies can strongly disrupt the quality of the maps (see **Figure 4f**)

**Supplementary Figure 7.**
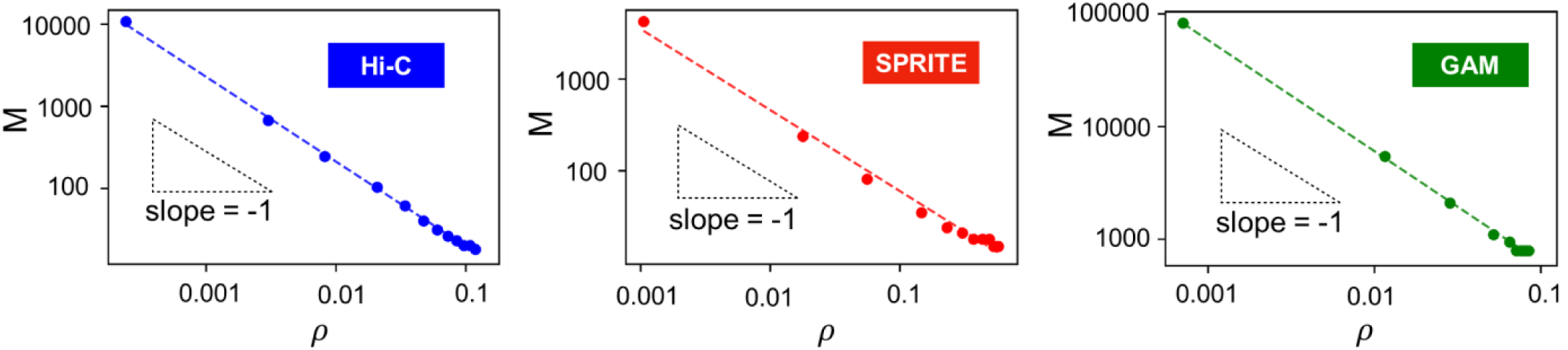
The estimated value of M is consistent with the Central-Limit-Theorem. In the *Sox9* locus case study, the values of M (see **Main Text, Figure 4**) at different efficiencies are plotted against *ρ*, the squared signal-to-noise ratio averaged over all the entries of a single-cell contact map (**Materials and Methods**). This is done for Hi-C (left), SPRITE (middle) and GAM (right). In all three plots (in log-log scale) the trend of M vs *ρ* is well fitted by a linear relationship with slope −1 (dashed lines) as expected by arguments based on the Central Limit Theorem (**Materials and Methods**).

**Supplementary Figure 8.**
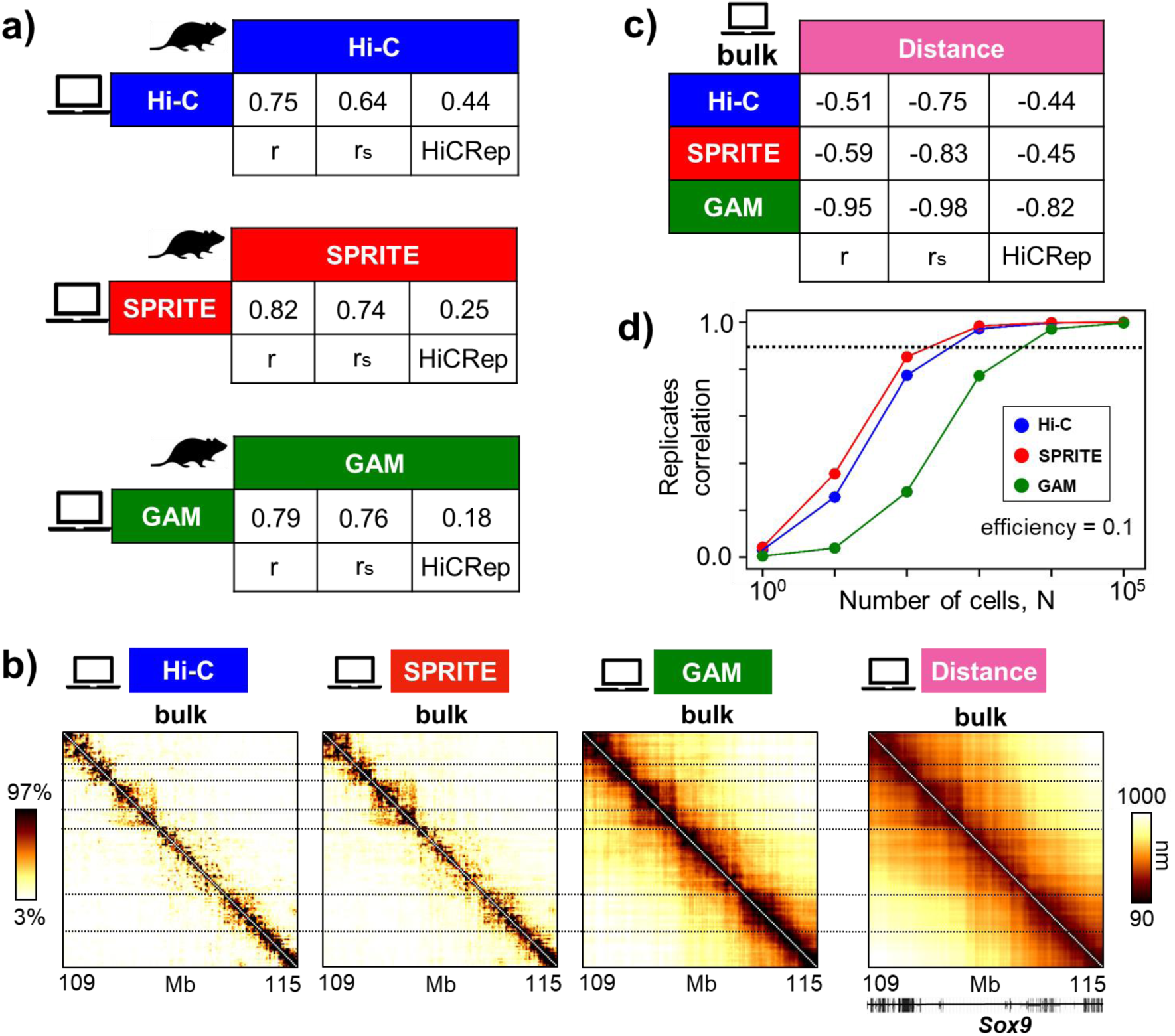
*In-silico* contact maps from 3D conformations derived from GAM data. a) We considered 3D structures for the mESC *Sox9* locus derived from GAM data^45^ (**Materials and Methods**). The corresponding Hi-C, SPRITE and GAM *in-silico* contact maps are compared to the experimental data (the same used in **Figure 1**, see **Materials and Methods**) and their Pearson, Spearman and HiCRep correlations are reported. b) The *in-silico* bulk contact maps are compatible with the average distance pattern obtained from the ensemble of GAM-derived 3D conformations. Horizontal lines are drawn to highlight the patterns detected across the contact and the distance maps. For the contact maps, color scale indicates the percentiles. c) Pearson, Spearman and HiCRep correlations are reported between each bulk contact map and the average distance map. d) Pearson correlation between replicates against the number of cells, for efficiency 0.1. The dashed line is the threshold correlation value r_t_=0.9. The values of M in this case are analogous to the ones obtained for the *Sox9, HoxD* and *Epha4* ensembles derived from Hi-C data, at the same 0.1 efficiency value (**Figure 4d, Supplementary Figure 2c** and **Supplementary Figure 3d**, respectively).

**Supplementary Figure 9.**
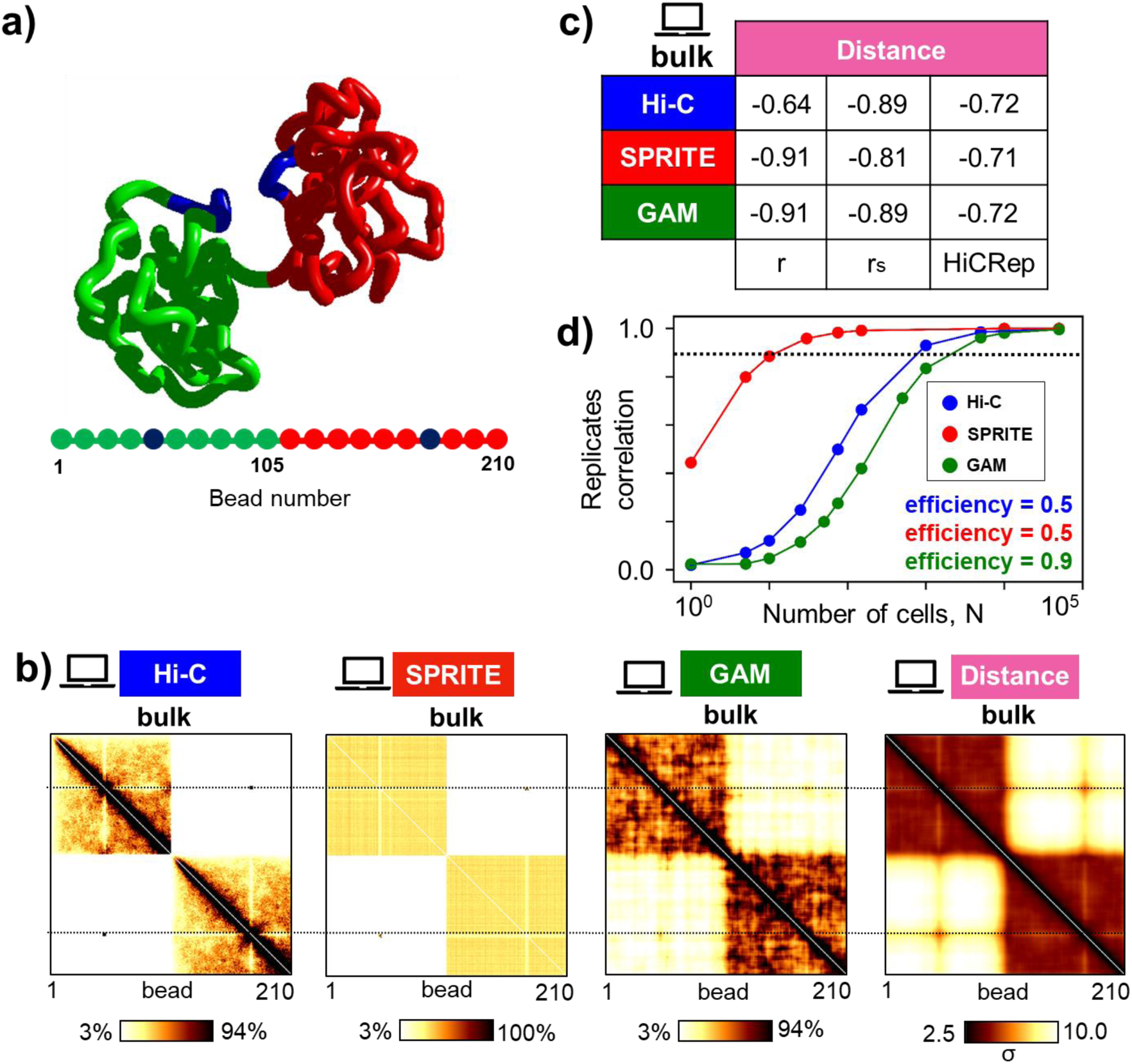
*In-silico* contact maps from 3D conformations of a toy polymer model. a) We considered a simple block copolymer model made of 210 beads where same colored regions attract each other (see **Materials and Methods**). The example of a 3D structure is shown. b) *In-silico* Hi-C, SPRITE and GAM bulk contact maps all yield contact patterns compatible with the average distance pattern derived from our ensemble of conformations. The horizontal lines are a guide to the eye. Color scale for the contact maps indicates the percentiles. For the distance map, color scale is given in the units of *σ*, the diameter of a polymer bead (**Materials and Methods**). c) The Pearson, Spearman and HiCRep correlations between each bulk contact and the average distance map are reported. d) Replicate Pearson correlations are plotted v.s. the number of cells, N, for: an efficiency equal to 0.5 for Hi-C and SPRITE, 0.9 for GAM. The values of M for the toy model are comparable to the ones obtained from the models of the *Sox9, HoxD* and *Epha4* loci at similar efficiencies (**Figures 4d,e**; **Supplementary Figure 2c, Supplementary Figure 3d, Supplementary Figure 8d**).

**Supplementary Figure 10.**
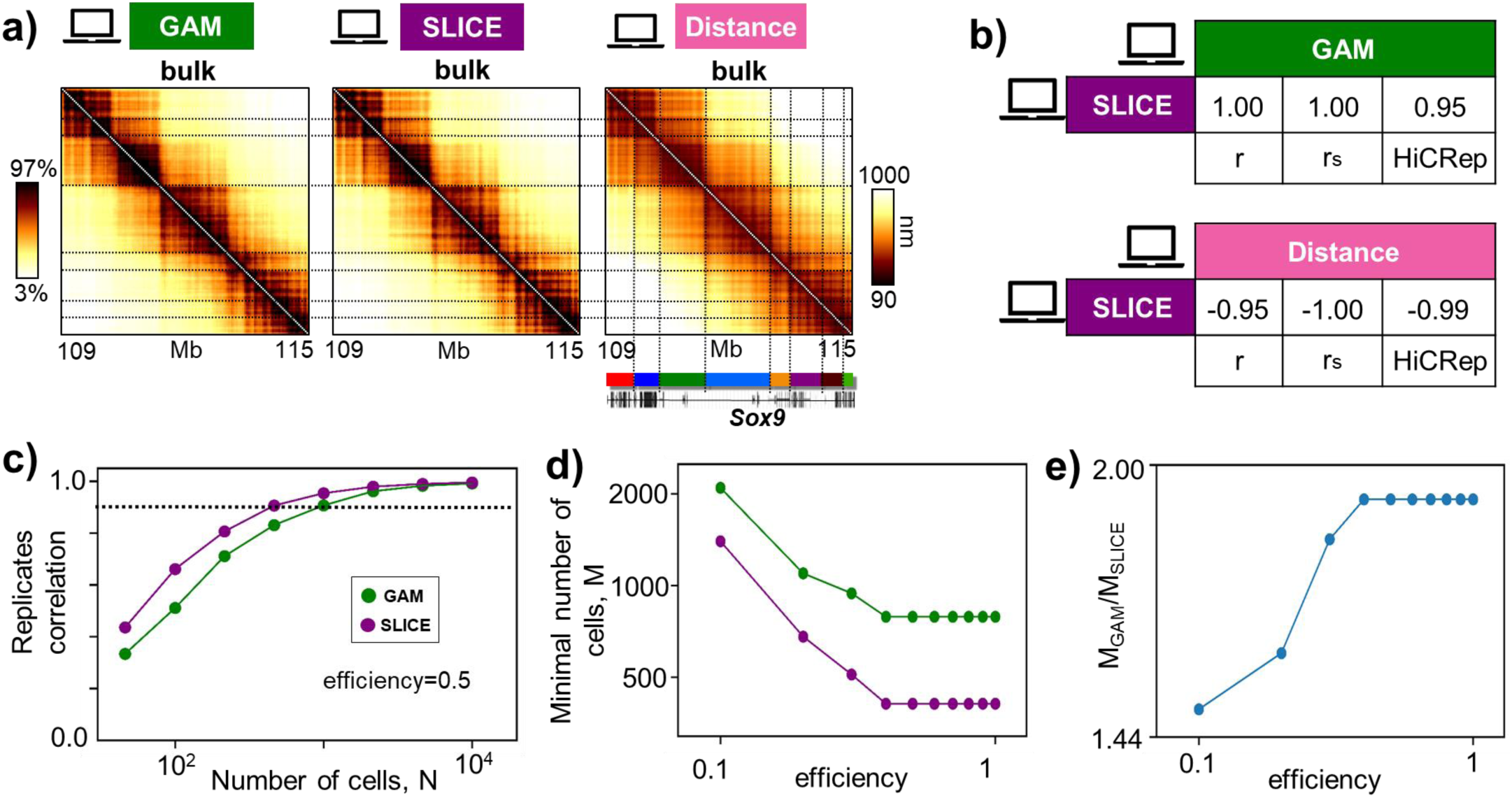
The SLICE analysis tool for GAM is faithful to the benchmark distance map. a) For the *Sox9* locus case study, the *in-silico* bulk GAM map and the corresponding SLICE map (**Main Text**) return consistent interaction patterns. Color scale indicates the percentiles of the maps. In particular, the SLICE single-cell interaction probability map is also faithful to the average distance pattern. Horizontal lines highlight that GAM and SLICE both capture the domain structure of the distance map, corresponding to the TADs^23^ shown in the color bar at the bottom. b) Pearson, Spearman and HiCRep correlations between SLICE and GAM bulk maps (top) and between SLICE and the average distance maps (bottom). c) The Pearson correlation between replicate contact maps is shown as a function of the number of cells, N, for GAM and SLICE at efficiency 0.5 (**Materials and Methods**). d) The minimal number of cells, M, to have reproducible replicates at different efficiencies for SLICE and GAM (as in **Figure 4f**). e) The ratio between M for GAM and for SLICE v.s. the efficiency. For efficiencies close to the experimental ones, say 0.5 or above^20^, the value of M for SLICE is approximately a factor two lower than for GAM.

## SUPPLEMENTARY TABLE

**Supplementary Table 1.**
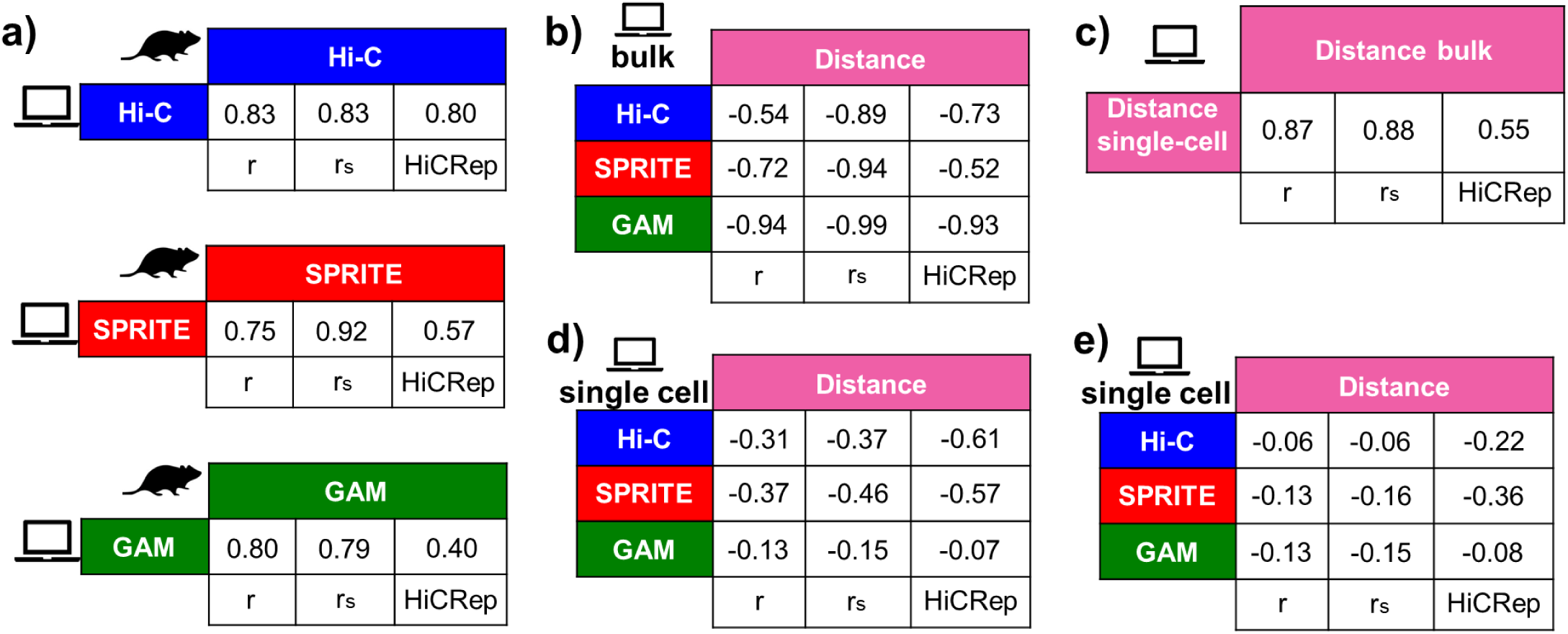
Pearson, Spearman and HiCRep correlations for the *Sox9* locus. a) Pearson (r), Spearman (r_s_) and HiCRep (scc) correlations between bulk Hi-C, SPRITE and GAM (from 1122 slices) experimental maps and the corresponding *in-silico* maps (see **Figure 1** and **Materials and Methods**), for the *Sox9* locus. b) Pearson, Spearman and HiCRep correlations between the average distance map and the *in-silico* bulk Hi-C, SPRITE and GAM contact maps (see **Figure 2**) for the *Sox9* locus. c) Pearson, Spearman and HiCRep mean correlations between *in-silico* single-cell distance maps and the average distance map for the *Sox9* locus (see **Figures 3a,b**). d) Mean correlations between *in-silico* single-cell contact maps - at efficiency 1 - and the corresponding single-cell distance map (see **Figures 3c,d**). e) Same as panel d), but here *in-silico* contact maps are generated with efficiencies similar to the experimental ones (0.05 for Hi-C and SPRITE and 0.5 for GAM; see **Main Text** and **Materials and Methods**). The reduction of efficiency worsens the similarity with the single-cell distance pattern.

## Notes

### Competing Interest Statement

The authors have declared no competing interest.

